# CapsidMesh: atomic-detail structured mesh representation of icosahedral viral capsids and the study of their mechanical properties

**DOI:** 10.1101/221663

**Authors:** José Luis Alonzo-Velázquez, Salvador Botello-Rionda, Rafael Herrera-Guzmán, Mauricio Carrillo-Tripp

**Affiliations:** Ciencias de la Computación, Centro de Investigación en Matemáticas, A.C., Jalisco S/N, Col. Valenciana, C.P. 36023, Guanajuato, Guanajuato, México; Laboratorio de la Diversidad Biomolecular, Centro de Investigación y de Estudios Avanzados Unidad Monterrey, Vía del Conocimiento 201, Parque PIIT, C.P. 66600, Apodaca, Nuevo León, México

## Abstract

Viruses are the most abundant pathogens affecting all forms of life. A major component of a virus is a protein shell, known as the viral capsid, that encapsulates the genomic material. The fundamental functions of the capsid are to protect and transport the viral genome, and recognize the host cell. Descriptions of this macromolecular complex have been proposed at different scales of approximation. Here, we introduce a methodology to generate a structured volumetric mesh of icosahedral viral capsids (CapsidMesh) based on the atomic positions of their constituents. Material properties of the capsid proteins can be set on every mesh element individually. Hence, we have control over all levels of protein structure (atoms, amino acids, subunits, oligomers,capsid). The CapsidMesh models are suitable for numerical simulations and analysis of a physical process using a third-party package. In particular, we used our methodology to generate a CapsidMesh of several capsids previously characterized by Atomic Force Microscopy experiments, and then simulated the mechanical nanoindentation through the Finite Element Method. By fitting to the experimental linear elastic response, we estimated the elastic modulus and mechanical stresses produced on the capsids. Our results show that the atomic detail of the CapsidMesh is sufficient to reproduce anisotropic properties of the particle.

## INTRODUCTION

Viruses are macro-molecular complexes made of nucleo-protein components. In the case of icosahedral viruses, at least 60 copies of a specific protein type (virus capsid protein, or VCP) self-assemble into a symmetric closed structure in the form of a hollow shell (capsid) encapsulating the viral genome (DNA or RNA). ^1^ Icosahedral capsids display interesting structural features, having a particular geometrical arrangement with a diverse range of VCP folds and particle size. The way proteins are arranged in the capsid is commonly described in terms of the so-called T-number.^2^ In recent years, X-ray crystallography and nuclear magnetic resonance (NMR) experiments have produced a considerable amount of structural information with atomic detail. ^3^

The study of the physical properties that characterize the viral capsid have advanced our knowledge in the fields of structural biology, biotechnology and medicine. In particular, virus nanoindentation produced by Atomic Force Microscopy (AFM) has been used as the standard way to analyze the particle’s mechanical response.^4^ AFM experiments have provided insights into the capsid’s molecular structure. ^5^ From a theoretical standpoint, it has also become possible to study whole viruses by molecular dynamics (MD) methods employing all-atom models.^6–9^ In particular, the nanoindentation experiments performed on the *Cowpea Chlorotic Mottle Virus* ^10^ have been computationally studied in some depth using coarse-grain models, ^11^ elastic network approximations, ^12^ and topology-based self-organized polymer models.^13^

Theoretical methods usually provide a higher space resolution than current experimental techniques, yet they tend to have an elevated computational cost. However, the theoretical framework of the Finite Element Method (FEM) is a promising and less expensive alternative approach. An important consideration is that FEM requires a mesh with particular regularity characteristics. Algorithms to generate a biomolecular surface mesh have been previously reported.^14–16^ In general, those meshes are highly optimized for molecular visualization,but are not robust enough for scientific numerical simulation analyses. In the case of viral capsids, some procedures first generate internal and external smooth surface meshes, ^17^ and then generate a non-structured volumetric model by filling the space found between them. Other methods do not use meshes but a set of kernel shape functions in an attempt to connect the atomic and continuum descriptions of mechanical properties.^18^

In this work, we present a methodology to build accurate structured volumetric meshes of icosahedral viral capsids based on atomic structural data, taking advantage of the inherent particle symmetries. Our approach has several benefits. 1) It can be systematically used to study the full icosahedral capsid diversity observed in nature ^3^ under the same framework. Furthermore, since it is geometrically conceived, it can be easily modified to analyze helical capsids as well. 2) Our meshes capture the spatial and geometrical arrangement of proteins in the capsid in great detail, i. e., the space between atoms is not artificially filled up, maintaining a “porous medium”. 3) It is possible to identify and extract the mesh elements associated to any set of atoms, residues or individual proteins, before and after numerical simulations. 4) Specific material parameter values can be set at each mesh element, independent of the structural level of the protein complex, allowing to test any scenario of material interfaces at different structural levels. Finally, to test our methodology, we carried out FEM simulations of the nanoindentation of several viruses, previously characterized with AFM, estimating the elastic modulus and mechanical stresses of the corresponding capsids with atomic detail. We show that our description is able to reproduce anisotropic properties of the particles.

## METHODOLOGY

To reduce the computational cost of the mesh generation procedure, we take advantage of the icosahedral symmetry of the virus particle by first meshing a subset of capsid proteins (GMesh). We then rotate images of this subset to generate the mesh of the entire capsid (CapsidMesh). The initial subset of capsid proteins for any icosahedral capsid is the so-called Icosahedral Asymmetric Unit (IAU). ^19^ The atomic structural information is specified in a file with the standardized coordinates of the IAU in PDB format. ^20^ A readily formatted data file can be downloaded from the *Infopage/Biodata* section of the virus of interest at the VIPERdb Science Gateway^3^ (Fig. S1). All the DNA or RNA structural information found in the data is discarded. Firstly, we consider the capsid to be enclosed by the surface of two concentric spheres connected through a radial tessellation projection. Then, we build several concentric intermediate spheres to generate hexahedral volumetric elements, discarding those that do not intersect any atom in the IAU. Images of the surviving hexahedrons are rotated and merged to produce the final structured volumetric mesh of the full capsid. The tessellation resolution can be adjusted to produce high definition meshes. We provide a detailed description of the steps involved in our methodology.

### Fundamental geometric unit (FGU)

Given an icosahedron and a sphere centered at the origin, one can project the edges of the icosahedron radially onto the surface of the sphere, producing an icosahedral tessellation with geodesic triangles (Fig. 1a). Any of the triangular faces of the icosahedron can be used to build the entire icosahedron by applying twenty rotations. The same is true for the icosahedral tessellation of the sphere. Furthermore, we can carry out the same process on a thickened up sphere and obtain an icosahedral tessellation of the shell with radial triangular prisms. We define any of the triangular prisms to be the fundamental geometric unit (FGU). The use of the FGU concept prevents the formation of intersecting mesh elements when the rotations are made later in the process, which is a typical problem found in other methodologies.

**Figure 1:**
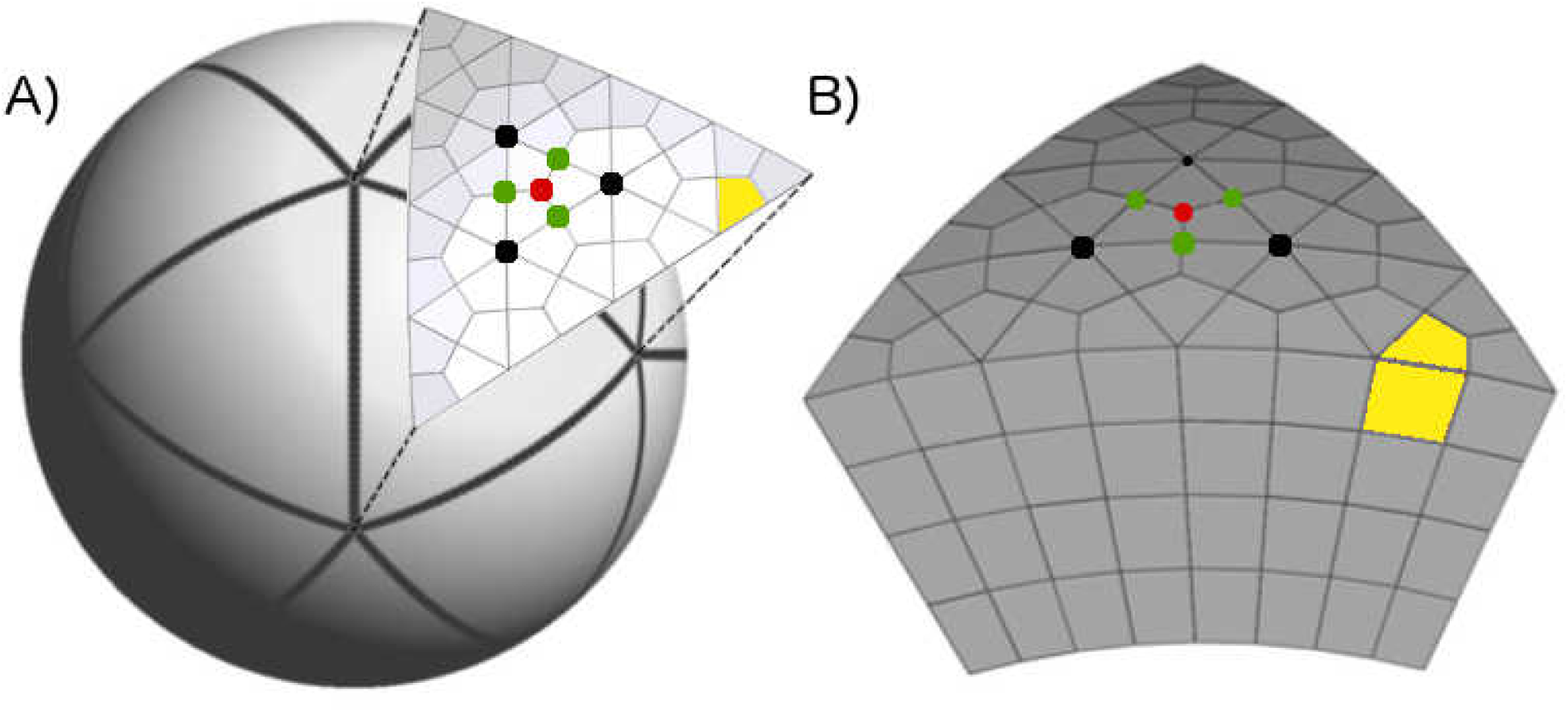
Construction of the fundamental geometric unit (FGU) mesh. (A) Icosahedral tessellation of a sphere with geodesic triangles. One face has been meshed with structured triangles. In this example, the vertices of one of the sixteen triangles on this face are depicted by black dots. Each triangle is subdivided into three non-square quadrilaterals using the baricenter (red dot) and the midpoint of the three edges (gren dots). The result is the 2D structured quadrilateral mesh. (B) Concentric spheres give layers of equal thickness. A radial projection of the face mesh produces structured hexahedrons, or mesh elements (one is highlighted in yellow). This is the 3D structured hexahedral mesh of the FGU.

### Mesh of the FGU

For simplicity, we first consider a thin FGU. We take one geodesic triangle of the tesellated sphere and give it a structured triangular mesh. We subdivide each equilateral triangle into three non-square quadrilaterals using the barycenter and the midpoint of the three edges. We call this grid the 2D structured-quadrilateral mesh. We project the line segments onto the spherical faces of the FGU. Each quadrilateral of the aforementioned 2D mesh produces two curved quadrilaterals on the curved faces of the FGU, that together make up the top and bottom faces of curved hexahedrons. In order to mesh a thick FGU that contains all the capsid’s atoms, we consider it has several layers of equal thickness given by thin FGUs, always using the same structured quadrilateral mesh. The result is the 3D structured hexahedral mesh of the FGU (Fig. 1b). In the example, the FGU’s mesh contains 16 *×* 3 *×* 4 = 192 hexahedrons, or mesh elements. An increase in mesh resolution will produce smaller hexahedrons and increase the number of mesh elements.

### Geometric icosahedral asymmetric unit (GIAU)

As mentioned before, we use the atomic coordinates of the capsid’s IAU as the starting point. To consider all the atom interactions, we construct a capsomere composed of all the adjacent asymmetric units of the IAU (Fig. 2a). In the example shown, we use a T=3 viral capsid. In this case, the IAU contains three identical capsid proteins occupying distinct interaction environments, identified with different labels and colors (according to the standard convention: A in blue, B in red, and C in green). Hence, the final capsomere is composed of 12 equivalent asymmetric units, including the central IAU, with 36 VCPs. We have to do this because we have to account for all the protein-protein contacts inside the IAU, and therefore, the FGU. The thickness of the FGU is defined according to the position of the closest and the farthest atoms from the capsid’s geometrical center found in the IAU (plus a threshold value). Therefore, all the atoms found inside the FGU, regardless of what asymmetric unit they belong to, are part of what we call the Geometric Icosahedral Asymmetric Unit, or GIAU (Fig. 2b).

**Figure 2:**
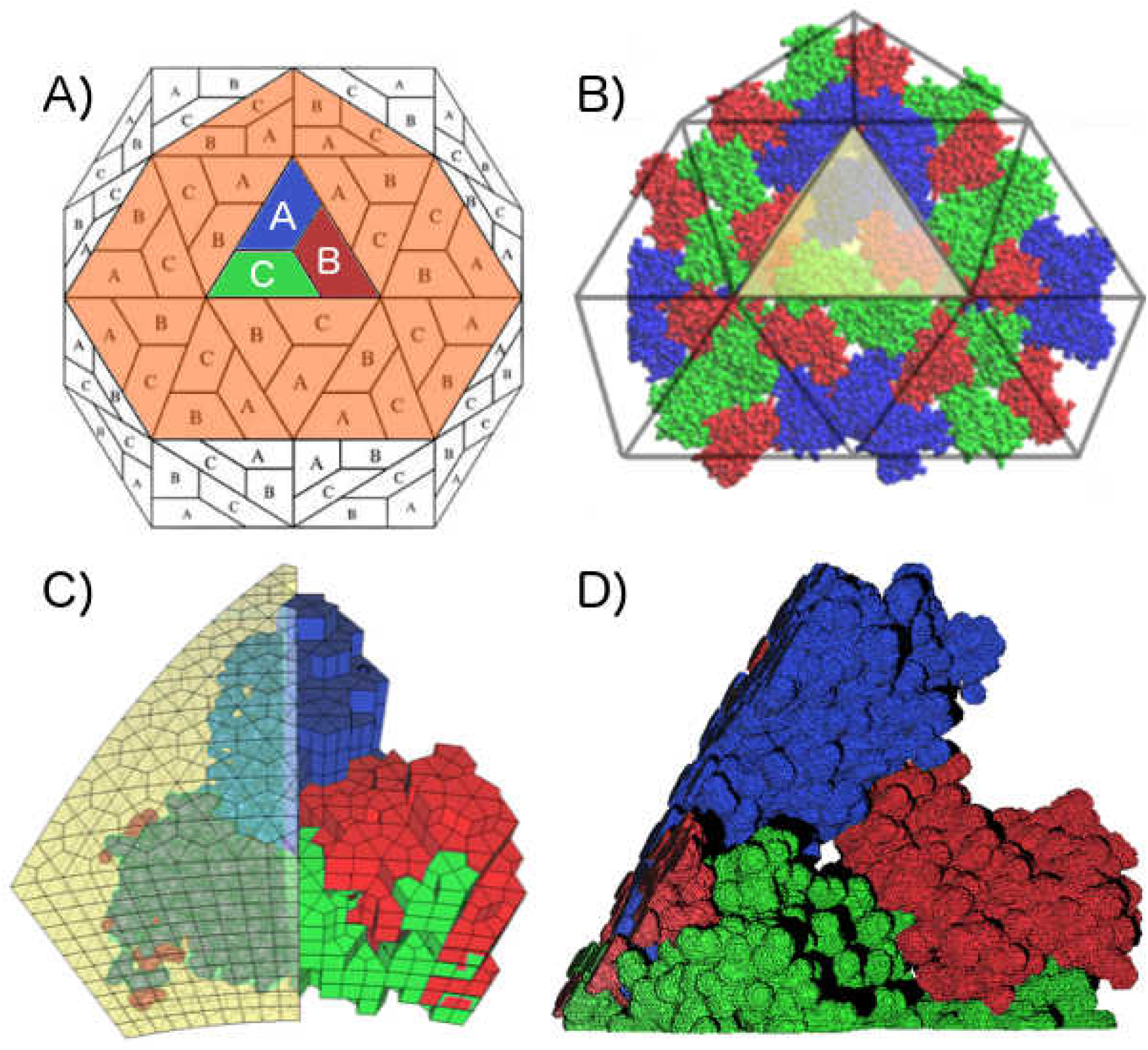
GMesh: construction and voxelization of the Geometric Icosahedral Asymmetric Unit (GIAU). (A) Schematic representation of a full T=3 icosahedral lattice. Each trapezoid represents an independent capsid protein occupying distinct environments. In this case, the central IAU is composed of three proteins, labeled A (blue), B (red), and C (green). All the adjacent asymmetric units are highlighted in orange. (B) Capsomere composed of 12 asymmetric units, 36 capsid proteins. Equivalent environments are depicted by color: A in blue, B in red, and C in green. (C) All the elements in the FGU mesh that intersect an atom in the GIAU are kept, otherwise discarded. An FGU mesh of 5Å resolution is shown. (D) The final set of mesh elements compose the GMesh. A resolution increase to 1Å produces an atom-detailed mesh.

### Radial voxelization of the GIAU

Next, we find all the elements of the structured hexahedral FGU mesh that intersect at least one atom in the GIAU according to the criteria explained in the next section. We discard those mesh elements that do not comply with these criteria (Fig. 2c). The mesh resolution is a parameter that is set in advance. When its value is high, a mesh with atomic detail is achieved. However, the number of elements increases drastically. The final set of mesh elements that are kept will be called GMesh (Fig. 2d).

### Intersection between FGU and GIAU

The volume of an atom of a given type is defined by the characteristic value of its radius. There are several radius definitions.^21^ Hence, we consider the atom type radii as parameters whose values can be set in a configuration file. Here, we use the van der Waals radius, defined as half of the internuclear separation of two non-bonded atoms of the same element type on their closest possible approach. We chose this definition because we found it gives better numerical stability in the simulations compared to other options (e. g., solvent accessible or excluded surface).

The criteria used to decide whether a hexahedral mesh element intersects a capsid’s atom are described in the pseudocode of algorithm 1 and Fig. S2. Several tests are made, where every FGU mesh element *e*_*i*_ is compared to every atom *p*_*j*_ in the GIAU. The variables involved are the atom radius r_*j*_, the distance d_*ij*_ between the center of the atom O_*j*_ and the barycenter of the mesh element G_*i*_, and the longest distance **a**_*i*_ between the element’s barycenter to a vertex A_*i*_.

#### Algorithm 1 Find mesh elements intersecting the volume defined by the van der Waals atom radius

~~~
**Require**: For *i* mesh elements and *j* atoms: array of FGU elements e_*i*_, array of GIAU atoms p_*j*_, and array with van deer Waals radii values r_*j*_.
1: **for** each *e*_*i*_ ∈ {FGU} **do**
2:      compute barycenter *G*_*i*_
3:      **for** each *p*_*j*_ ∈ {GIAU}g **do**
4:            compute distance *d*_*ij*_ between *G*_*i*_ and the center *O*_*j*_ of *p*_*j*_
5:            **if** *d*_*ij*_ < r_*j*_ **then**
6:                  RETURN: element *e*_*i*_ is kept
7:            **else**
8:                  compute max(*a*_*i*_), the distance between *G*_*i*_ to all vertices of element *e*_*i*_. The farthest vertex from *G*_*i*_ is labeled *A*_*i*_
9:            **end if**
10:            **if** d_*ij*_ < (*r*_*j*_ + *a*_*i*_) **then**
11:            **if** vertex of element *e*_*i*_ is inside of atom *p*_*j*_ **then**
12:                  RETURN: element *e*_i_ is kept
13:            **else if** Oj is inside *e*_*i*_ **then**
14:                  RETURN: element *e*_*i*_ is kept
15:            **end if**
16:      **end if**
17: **end for**
18: **end for**
~~~

### CapsidMesh: mesh of the full capsid

Once the voxelization of the GIAU has been produced, we make images and transform their positions to generate a complete icosahedral particle, according to the set of VIPERdb’s standard rotation matrices. The process of merging the GMesh images consists of finding all element vertices occupying the same position in space. This is a computationally expensive task. A way to considerably reduce the number of comparisons is to identify which mesh elements are at the interfaces of the FGU and, therefore, the GMesh (Fig. 3a,b). Consequently, in the end we know which hexahedral elements belong to what atom, residue, protein or GMesh interface (Fig. 3c). At this point, the mesh of the full capsid, or CapsidMesh, is ready to be used in numerical simulations studies (Fig. 3d). The mesh resolution is a free parameter that controls the coarseness of the final model (Fig. S3).

**Figure 3:**
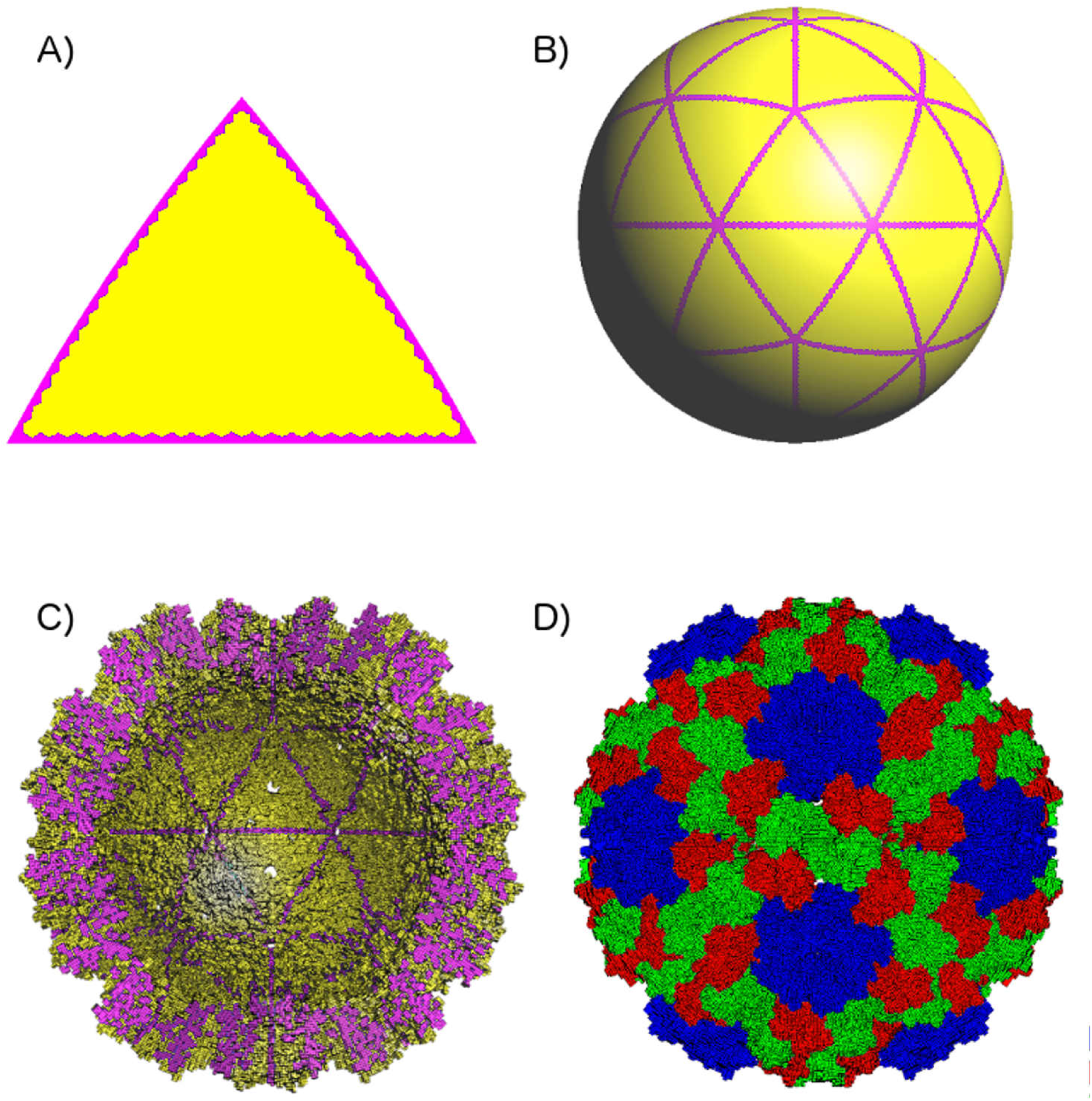
CapsidMesh of a T=3 viral capsid. (A) All the FGU mesh elements at the interfaces are identified, shown in purple. (B) Only those elements are used in the merging of the images process. (C) Inner view of the final volumetric mesh. The GMesh interfaces (purple cells) and the capsid’s atomic detail are clear. (D) The CapsidMesh is the final volumetric structured mesh. The hexahedral elements are colored according to the protein environment they belong to (A in blue, B in red, and C in green).

### Finite element analysis of CapsidMesh nanoindentation

Here we give a brief description of the theory behind the nanoindentation numerical sim-ulations using the Finite Element methodology. The relation between strain and stress is deduced from the elasticity property, generally expressed via a constitutive-rigidity matrix

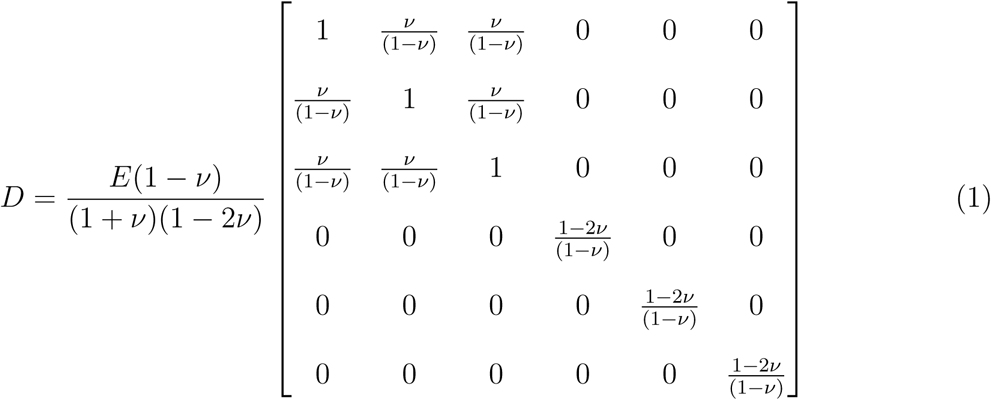

given by a symmetric 6 *×* 6 matrix which contains the elastic parameters of the medium. In the anisotropic case, it has two independent parameters. For isotopical and homogeneous materials, the two parameters involved in such matrix are the elasticity modulus *E* (or Young’s modulus) and the Poisson coefficient *ν*. Note that, for heterogeneous materials, the number of parameters involved may be as large as twenty one. Hooke’s law states that

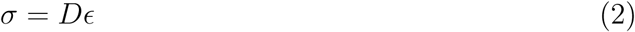

where *σ* is the stress and *ε* the strain. The deformation is modeled by the linear partial differential equation given by Newton’s second law:

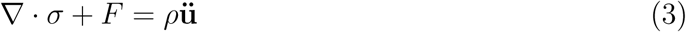

where **u** is the displacement, *F* is the force applied per unit of volume, and *ρ* is the mass density. We can describe the material internal displacements through a discretization of space. Such discretization is achieved by the CapsidMesh. Interpolation of the values of the variables on the nodes of the mesh create a field. Such interpolation is achieved via functions *N*_*i*_ of the form

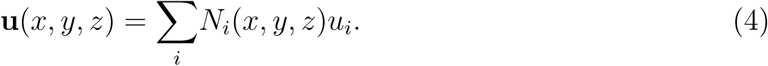

Multiplying Newton’s second law by some weight functions *W*_*i*_, which can be the same as the interpolation functions (Galerkin technique^22^), and integrating over the entire domain as well as neglecting the inertia forces, we get the following linear system:

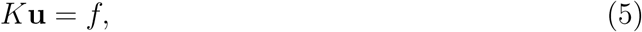

generated by the coupling of all the systems given by all the mesh elements, so that *K* is the standard rigidity matrix of FEM and *f* is the vector of forces.^22^

This equation must be solved numerically and efficiently with techniques such as LU factorization, iterative methods of preconditioned conjugate gradient, etc. Once the displacements have been obtained, the stresses can be computed at the points of integration by using Eq. 2. Such stresses can be easily extrapolated to the nodes of the mesh using smoothing techniques.^22^ One way to understand the effect of the stress tensor with the help of a scalar is to use the von Mises stress J2^22^

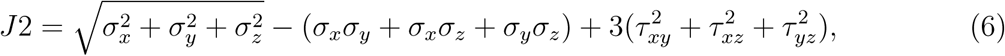

where *τ*_*xy*_, *τ*_*xz*_, *τ*_*yz*_ are the components of the shear stress tensor.

In this way, one can quantify and visualize the effects of the degree of stress in each point of the domain of the mesh. This is an important result since all materials have an equivalent resistance stress with which one can determine how close or how far from failure is each point of the mesh depending on the deformation force imposed on the capsid.

### RESULTS

### Mesh generation

We used our methodology to generate a CapsidMesh of icosahedral capsids based on the atomic Cartesian coordinates of the IAU found in VIPERdb (Fig. S1). A previously reported methodology to build low-resolution density maps and triangulated isosurface smoothed meshes, tessellated with tetrahedra, also based on the atomic coordinates of a capsid has been used in a series of studies. ^23–25^ In particular, a comparison between the *Cowpea Chlorotic Mottle Virus* volumetric mesh generated with such methodology and a mesh generated with the CapsidMesh methodology is shown in Figure 4. Clear differences can be seen in the internal structure regularity and the surface detail of the meshes. The mesh generated with our methodology is structured and reproduces the molecular surface with atomic detail, both on the capsid’s interior and exterior, as well as the spaces between protein interfaces. In addition, all the elements in the mesh can be pinpointed and assigned to any structural feature of the particle (Fig. S4). To test the robustness of our method, we generated a mesh for a set of representative viral capsids on a wide range of T-numbers: T1, T2, T3, T4, T7d, T7l, T13, T27, and T31 (Fig. S5). Our methodology was capable to systematically generate a mesh for a large range of capsid sizes available on VIPERdb (approximately from 200°ÅA to 1000°ÅA in diameter). Even more, it was possible to generate a mesh with the correct shape for capsids with known problematic structural features, such as constitutive holes or large protrusions (e. g. 3kk5 or 3j31, respectively. 4g93 is an example of a case presenting both features).

**Figure 4:**
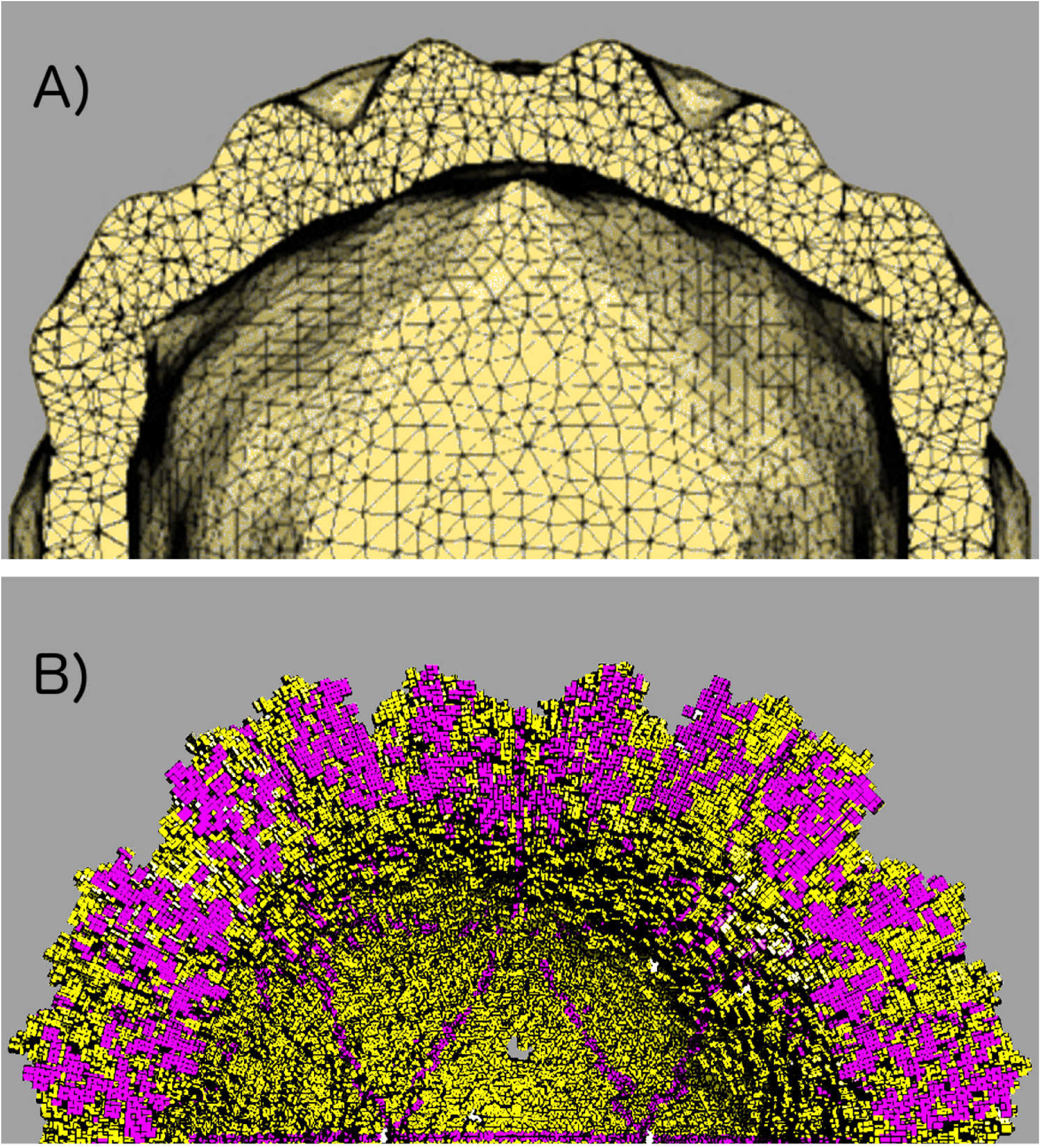
*Cowpea Chlorotic Mottle Virus* capsid. Zoomed in cut view of the 3D mesh created with the (A) method reported by Gibbons and Klug, ^24^ and (B) CapsidMesh method reported in this work. Differences in mesh structure regularity are appreciated. GMesh interfaces in purple.

### Nanoindentation numerical simulations

The AFM experiment measures the relation between the applied force and the capsid indentation, i. e., a quantification of the particle’s deformation response to squeezing. The experiment records a characteristic force-indentation profile (experimental FZ curve), which is used to estimate the value of the capsid’s mechanical properties by fitting a theoretical model to the collected data. The slope of the FZ curve in the linear regime is related to the capsid’s spring constant *k*.^26^ AFM experimental results have shown that deformations as high as 30% of the particle’s radius still produce a reversible elastic linear response.^5^,26–29 It is hard to control the precise contact point where the cantilever touches the capsid in the AFM experiments. If the particle were a perfect sphere, this would not be of importance. However, icosahedral capsids have several symmetry axes, which in principle, might have a different mechanical response. In numerical simulations, it is straightforward to set a particular capsid orientation in line with the force load applied. In this way, nanoindentation simulations can be achieved on a specific symmetry axis.

In this work, we combined the CapsidMesh methodology with the FEMT tool^30^ to reproduce AFM capsid nanoindentation experimental results to test and characterize the generated meshes. The setup of the simulation is depicted in Figure 5. Once a symmetry axis was selected and the capsid mesh put in the right orientation, indentation on the capsid was produced by applying a controlled force load with a probe in direct contact with the top mesh elements. The capsid was supported by a base in direct contact with the bottom mesh elements. Both the probe and the base were nondeformable solids. Since there are no tangential forces, there is no sliding on the system. The indentation produced was quantified by the displacement measured in each mesh element.

**Figure 5:**
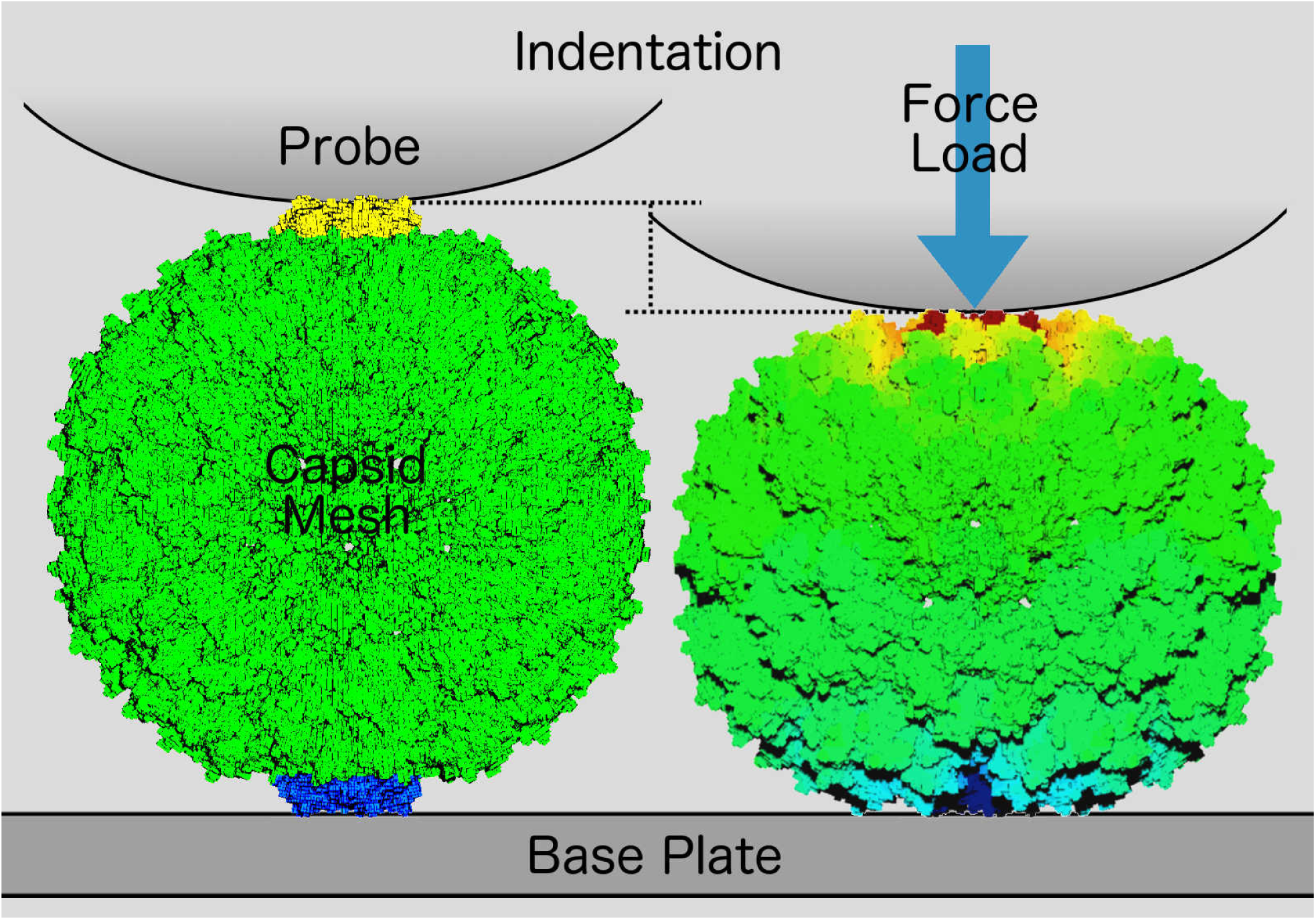
Capsid nanoindentation simulation. Left: Initial conditions, where the mesh elements in blue are in direct contact to a base, and mesh elements in yellow are in direct contact with a probe. Right: A force load is applied on a given capsid’s symmetry axis. The mesh deformation is quantified by the indentation produced (displacement measured in each mesh element). The color scale used represents minimum displacement in blue, and maximum displacement in red.

We performed FEM nanoindentation simulations of viruses that had been previously characterized through AFM and whose atomic coordinates were available, namely, *Cowpea Chlorotic Mottle Virus* (T=3),^26^ *Human Hepatitis B Virus* (T=4),^27^ *Bacteriophage T7 Prohead* (T=7),^28^ and the *HK97-like bacteriophage HII* (T=7)^29^ (Fig. 6a). The goal was to find the set of values for the parameters *E* and *v* such as to reproduce the experimental spring constant *k* for each capsid. It was previously shown that the simulation results are rather insensitive to the Poisson coefficient for values *v <*0.5.^26^ We corroborated that observation with our model and set *v* to a constant value of 0.3 to be consistent with previous reports. Using an initial guess for the force load and for the elastic modulus within the known linear regime range, capsid indentations on the 5-fold symmetry axis were recorded as a function of mesh resolution. Then, a linear scaling of the form *E*_1_*/E*_2_ = *I*_1_*/I*_2_ was applied to each point in order to find the elastic modulus value that gave the expected experimental force/indentation ratio *k* (Fig. 6b). Also, we conducted indentations on the 3 and 2-fold axes on a mesh with a resolution of 1Å (atomic detail), recording the force needed to produce the same indentation as in the 5-fold axis. Results are summarized in Table 1.

**Table 1:**
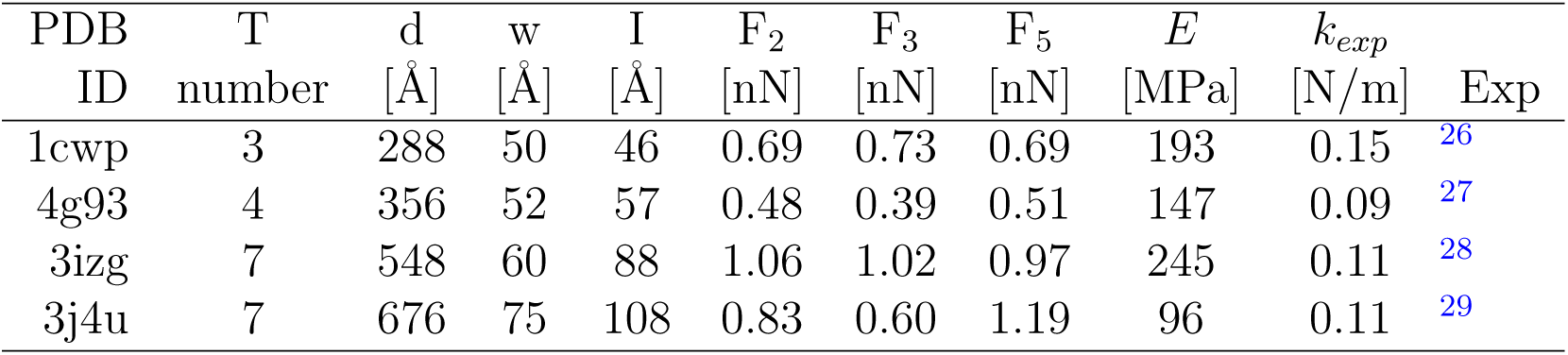
Nanoindentation of wild type empty capsids and mechanical properties of *Cowpea Chlorotic Mottle Virus* (1cwp), *Human Hepatitis B Virus* (4g93), *Bacteriophage T7 Prohead* (3izg), and *HK97-like bacteriophage HII* (3j4u), using a mesh resolution of 1Å in all cases. Shown: geometrical protein arrangement *T*, diameter *d* (two times the outer radius), capsid width *w* (outer radius minus inner radius), nanoindentation *I* (16% of *d* in all cases), force needed to produce *I* on the 2-fold, 3-fold, and 5-fold axes, Young’s modulus *E* prediction, experimental spring constant *k*_*exp*_, and data source.

**Figure 6:**
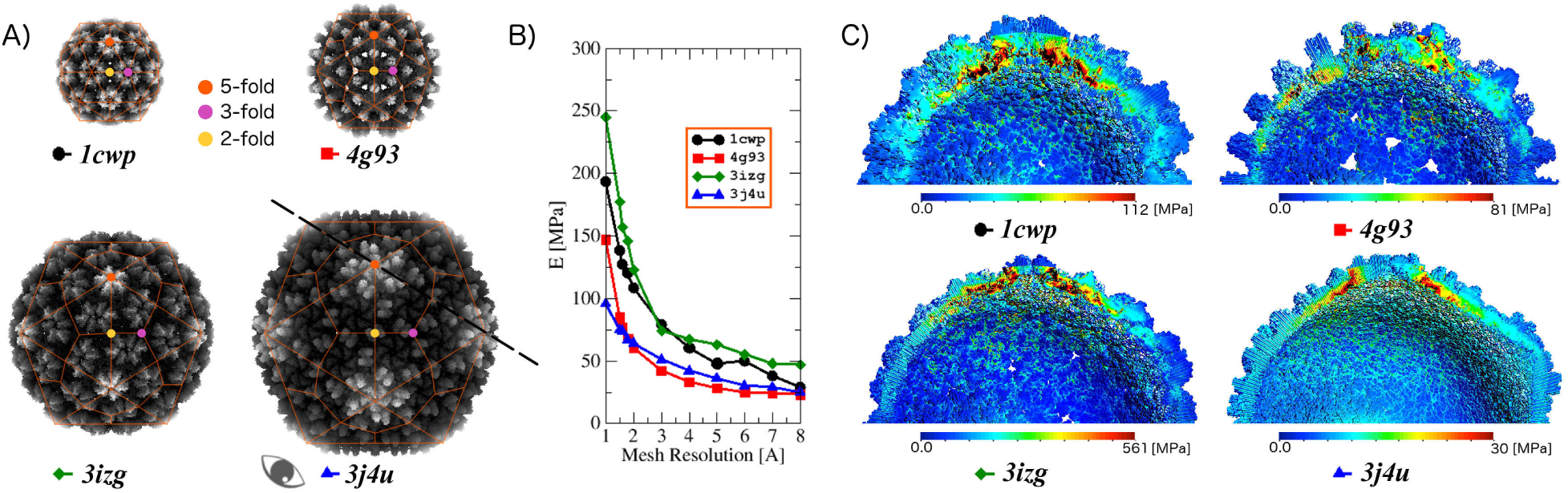
Characterization of CapsidMesh and mechanical properties. (A) The rendered molecular shape of *Cowpea Chlorotic Mottle Virus* (1cwp, black circle), *Human Hepatitis B Virus* (4g93, red square), *Bacteriophage T7 Prohead* (3izg, green diamond), and *HK97-like bacteriophage HII* (3j4u, blue triangle). The corresponding location of a 5-fold, 3-fold, and 2-fold axes are indicated. The viewpoint and plane cut used in panel C is shown. Capsids are proportionally sized, generated with VMD. (B) Dependence of Young’s modulus *E* on the mesh resolution to reproduce the experimental spring constant *k*. (C) Stress analysis after nanoindentation. Transverse cut showing the interior of the capsid. von Mises values are shown for a force load of 0.5 nN using the E value corresponding to a 1Å mesh resolution.

The von Mises stress analysis for the 5-fold indentation is shown in Figure 6c. It is clear where the regions of maximum stress are (around the loading point as expected), irrigating towards the equator line. Interestingly enough, we found that regions away from the loading point also present high von Mises stress values between VCPs, i. e., in the protein-protein interfaces. Also, it becomes clear how the mesh captures the true shape and atomic distribution of the capsids, e. g. the protrusions and holes of the *Human Hepatitis B Virus*, or the deviation from a sphere in the case of the larger *HK97-like bacteriophage HII* capsid. Such mesh fidelity to the molecular shape allows the observation of how the structural features affect the stress distribution both outside and inside the protein shell. Those are natural results of our model, evidence of the importance of having a faithful description of the molecular surface.

## DISCUSSION

There are several advantages of the CapsdiMesh methodology over previously reported methods to generate mesh models of icosahedral capsids. One is the level of control the user has over each element of the structured mesh. The volumetric meshes produced with our methodology are faithful to the spatial distribution of the atoms and the capsid’s VCPs geometrical arrangement, i.e. there are no mesh elements in regions corresponding to empty space between proteins. This is to be contrasted with the meshes produced by previous methods,g.,^17^ where the space between two surface meshes (capsid’s interior and exterior) are completely filled up. Our meshes have a radial structure built with regular elements and are suitable to be used in numerical analysis methods such as finite element, finite volume and finite differences. Moreover, one can assign specific material characteristics to each element in the mesh and, therefore, have control over the different structural levels of the macromolecular structure, e. g., atom, residue, oligomer. Our method converts the atomic models to meshes, whereas the simulation and analysis of the meshes are carried out by a third party software. In our case, we used FEMT, however, the way we designed the CapsidMesh methodology offers the flexibility to use other simulation and analysis packages. This independence increases the value of our methodology.

Given that even if a mesh element has a minor intersection with an atom the entire element is accepted into the mesh, the CapsidMesh is always an over-approximation of the actual volume occupancy. Also, the mesh produced does not have a smooth surface, therefore, the resulting mechanical simulations will likely produce stress concentrations at the point were the force load is applied. Both effects are greatly diminished by setting a high mesh resolution value. However, they should not be a problem when regularity in the mesh is a desirable property and it does not affect simulations of molecular properties that could be very sensitive to area or volume. We are working on the implementation of refinement techniques to address such problems when that is not the case. Likewise, the CapsidMesh methodology takes advantage of the intrinsic symmetry of icosahedral capsids to reduce the computational cost involved in the generation of a volumetric structured mesh. We focused our work on that type of capsid geometry because there is a large amount of structural data available. To the best of our knowledge, it is also the only geometry for which there are experimental AFM nanoindentation studies, as opposed to helical capsids. However, it is possible to adapt the procedure to other geometries, e. g. cylindrical symmetry, keeping the computational efficiency. With this in mind, cellular protein assemblies could also be meshed following our procedure by finding the correct symmetry. In the case of molecular complexes without symmetry, the meshing would have to be over a cubic domain and the computational cost should be decreased by other techniques. However, the mesh element-atom intersection algorithm can be the same as that in the CapsidMesh approach. We are also working on the development of such methodologies.

We produced the CapsidMesh of several viruses and performed a simulated nanoindentation on them through FEM. By reproducing experimental AFM measurements, we used our models and methodology to estimate the value of the Young’s modulus *E*, an important macroscopic material property. It is worth noting that *E* might not be the best descriptor to use for a microscopic capsid, since it is not an homogeneous medium and it has an anisotropic character. Nonetheless, several works have estimated the value of *E* for the *Cowpea Chlorotic Mottle Virus* in the past. Such estimations seem to highly depend on the approximation used in the continuum model fitted to the experimental data. The simplest model for the mechanical response is to think of the capsid as an elastic thin spherical shell undergoing small deformations. Using this approximation, *E* has a value of 140 MPa.^26^ A homogeneous, isotropic, elastic, thick-shell non-linear continuum elasticity model roughly captured the linear behavior observed in experimental measurements giving estimates for *E* of 280-360 MPa.^23^ A heterogeneous geometry, non-linear continuum elasticity model was tuned with a value for *E* of 215 MPa to match the experimental spring constant value. ^24^ Even more, the use of a mesh free method scaled to the experiments produced an effective *E* of 493 MPa.^18^ The last methodology is based on the atomic coordinates of the capsid, hence providing the same advantage as our method in terms of attempting to bridge the atomic and continuum descriptions.

Our model, being a continuum approximation, does not take into account long-range atomic interactions explicitly. On the other hand, short-range interactions are present by the inter-atomic overlap produced by the van der Waals radii. Atoms within a covalent bond distance will have a large overlap, whereas atoms within a van der Waals interaction distance range, e. g. VCP-VCP interface atoms, will have a smaller overlap but still be connected by elements of the mesh, even at the highest mesh resolution. However, one has to keep in mind that continuum models can only capture geometric details up to a lower limit, as previously shown.^31^ In this sense, the mesh resolution is a parameter that controls the coarseness of the model in our methodology. For mesh resolutions > 15Å, all cases studied in this work converge in terms of the elastic response *E*, indicating an upper limit on the coarseness and lack of detail on the model. As we increase the mesh resolution, structural differences between capsids start to gain relevance, up to a point where they can be clearly distinguishable from each other. We found this to be at 1Å of mesh resolution. Higher resolutions, even though possible, would not add much to the description of the continuum model. We believe such lower limit is a good balance between a fine geometric detail and a coarse-scale of deformations for our model. In this context, we present the relationship between the mesh resolution and the Young’s modulus value that reproduces the experimentally measured spring constant *k* for the four cases studied. The *E* values predicted by our model are in the same order of magnitude as those previously reported, when the mesh resolution is on the atomic level. Furthermore, within our approximations, the structural detail of a Capsidmesh is sufficient to obtain anisotropy. To the best of our knowledge, CapsidMesh is the only model that reproduces the correct relation between symmetry axes in terms of stiffness. AFM experiments showed that the icosahedral axes of T7 capsids follow the relationship 2 *>* 3 *>* 5.^28^ Our prediction for the *Bacteriophage T7 Prohead* is concomitant with that observation. This result is one more indication that the use of a mesh with atomic detail is important in the description of macromolecular complexes. With this in mind, and even though we only used CapsidMesh-FEMT to study the nanoindentation process, we expect that the estimation of other biophysical properties would also improve.

## CONCLUSIONS

In this work, we present a methodology to systematically generate atomic-detailed regular structured volumetric mesh representations of icosahedral capsids based on atomic data. With the use of a CapsidMesh, it is straightforward to assign a set of mesh elements to any given structural protein level, i. e., from individual atoms to a whole VCP. The CapsidMesh can be coupled to a third party simulation and analysis tool to study mechanical properties of the particle. In this case, we used FEMT to simulate the nanoindentation process of viral capsids previously characterized through AFM experiments. By reproducing the elastic response through the experimental spring constant, the CapsidMesh-FEMT combination is able to estimate the Young’s modulus, showing consistent results with previous findings on organic materials. In addition, our results show that a mesh with atomic-detail is sufficient to produce an anisotropic response of the icosahedral particles concomitant with experimental results.

## Acknowledgement

Simulations were executed in the high performance computing center Insurgente from CIMAT, administered by Jesús Rocha. We gratefully acknowledge MSc. Jorge Lopez-Ruiz from CIMAT for his technical assistance by conducting some of the simulations presented here, and the support of NVIDIA Corporation by the donation of the Titan X Pascal GPU used in this work. R.H.G. would like to thank the International Centre for Theoretical Physics and the Institut des Hautes Études Scientifiques for their hospitality and support.

## Supporting Information Available

## Supplemental information

**Figure S1.**
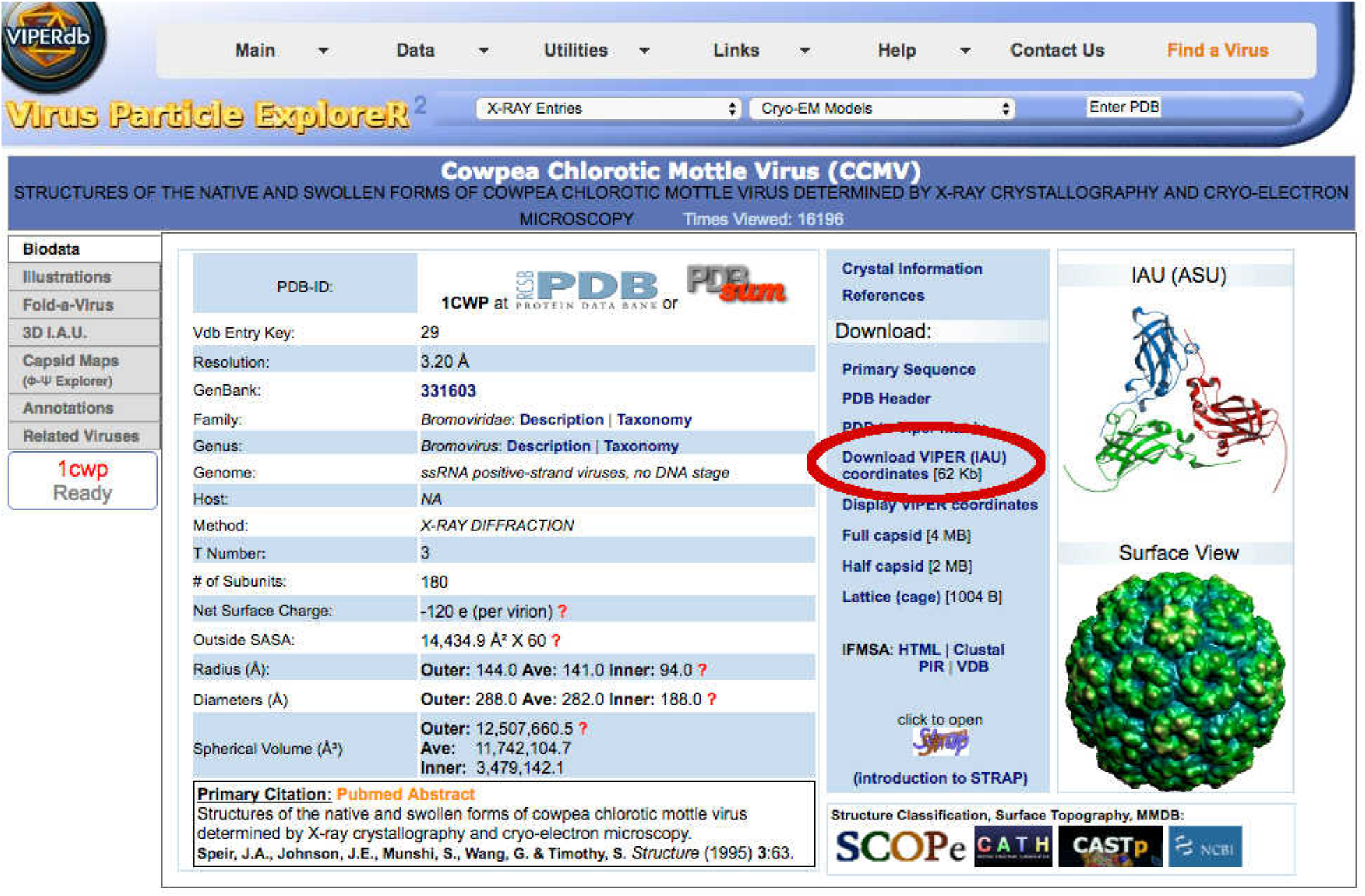
Infopage of a particular virus at the VIPERdb Science Gateway (http://viperdb.scripps.edu/). As an example, we show the data of the *Cowpea Chlorotic Mottle Virus* (PDBID:1CWP). The link to download the standardized atomic coordinates of the IAU is highlighted.

**Figure S2.**
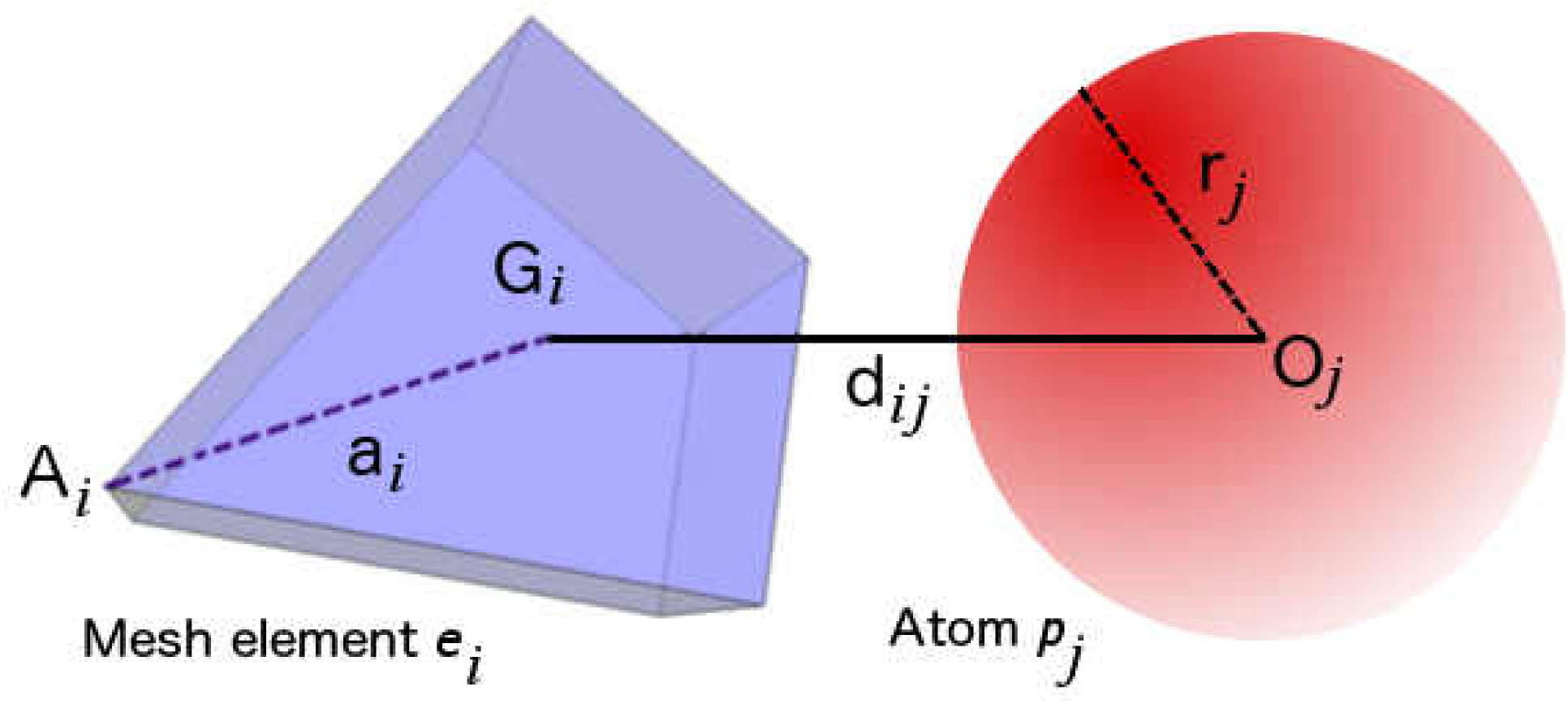
Intersection between FGU elements and GIAU atoms. We perform several tests, where every FGU mesh element *e*_*i*_ is compared to every atom *p*_*j*_ in the GIAU. The values involved in the tests are the atom radius r_*j*_, the distance d_*ij*_ between the atom’s center O_*j*_ and the element’s barycenter G_*i*_, and the longest distance **a**_*i*_ between the element’s barycenter to the vertex A_*i*_.

**Figure S3.**
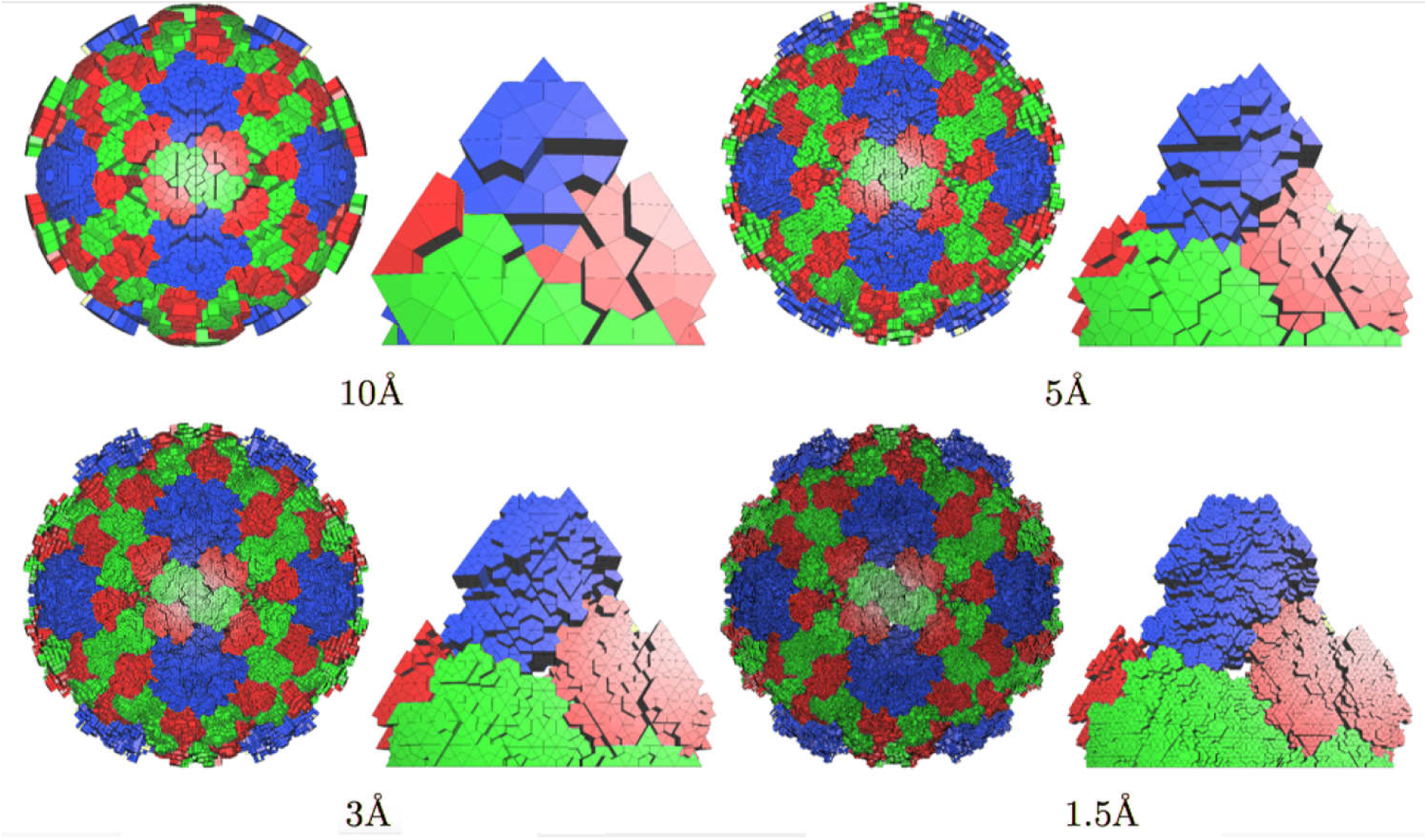
Coarseness of the mesh generated by CapsidMesh. The resolution is a parameter set by the user. Four different resolutions are shown for the same T=3 capsid. The full capsid and the corresponding GMesh are shown for each case. Mesh element color indicates distinct protein environments in the capsid: A in blue, B in red, and C in green.

**Figure S4.**
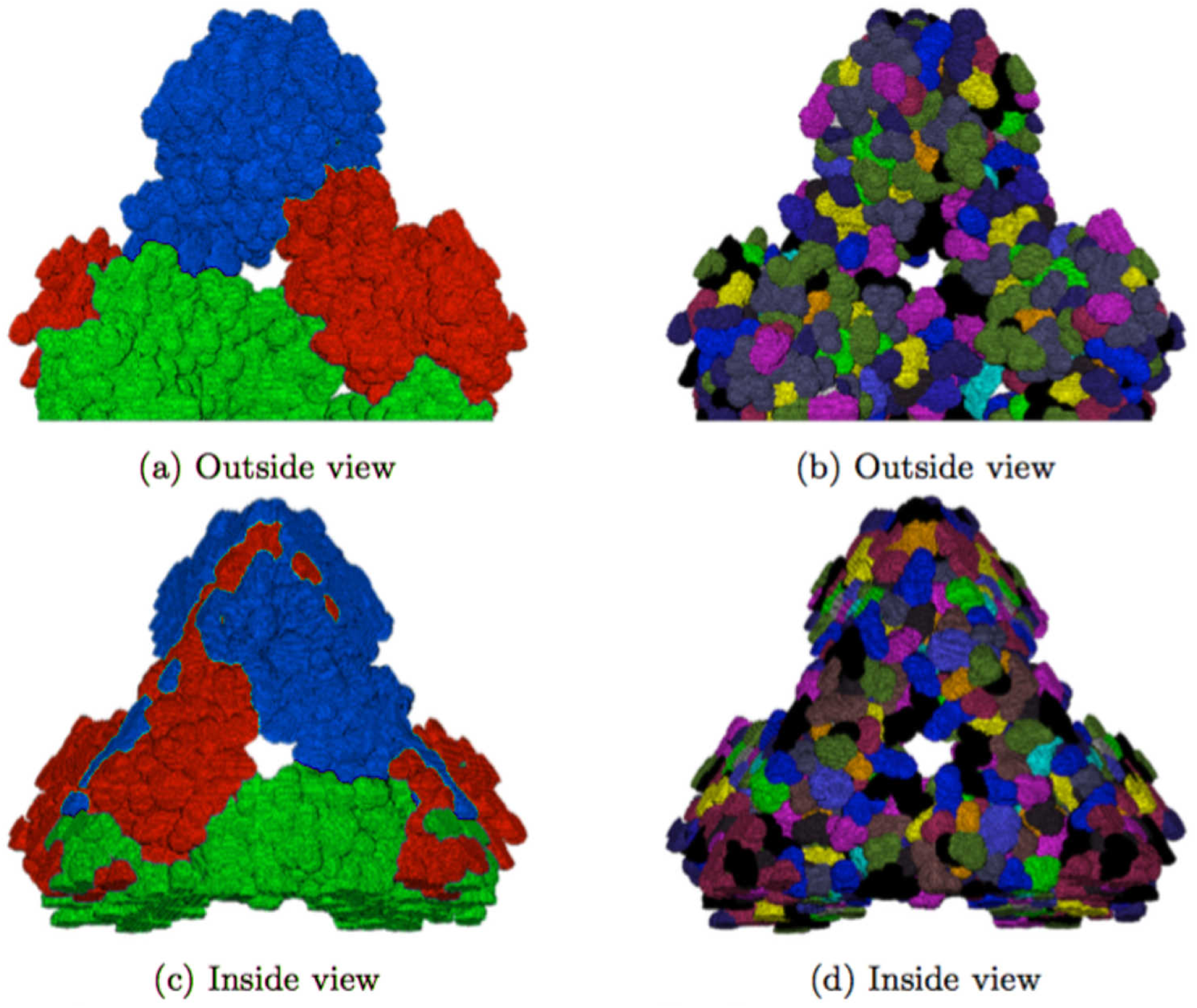
Assignment of specific properties to each element in the mesh. GMesh of the CCMV shown from the outside and from the inside of the capsid. On the left, mesh element color indicates distinct protein environments in the capsid: A in blue, B in red, and C in green. On the right: residues colored by amino acid type.

**Figure S5.**
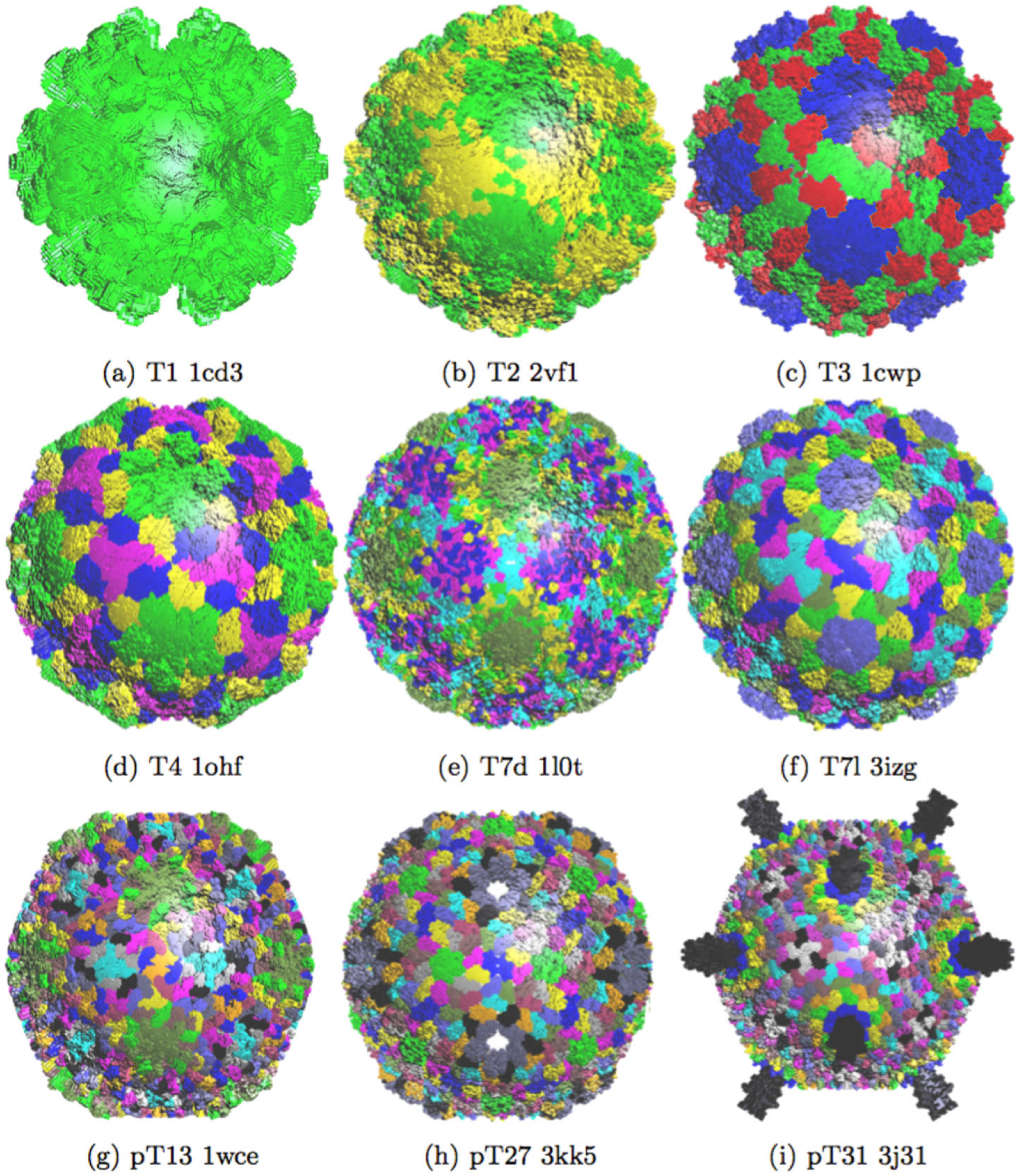
CapsidMesh of a representative set of virus capsids with different T-number. The color of each mesh element indicates the location of proteins in different environments inside the capsid. The PDBID and the T-number value is shown under the corresponding mesh.

## References

(1) Cann, J. A. Principles of molecular virology; Elsevier Ac. Press (4th ed.), 2005.

(2) Caspar, D. L. D.; Klug, A. Physical principles in the construction of regular viruses; Cold Spring Harbor Laboratory Press, 1962; pp 1–24.

(3) Carrillo-Tripp, M.; Shepherd, C. M.; Borelli, I. A.; Venkataraman, S.; Lander, G.; Natarajan, P.; Johnson, J. E.; Brooks, C. L.; Reddy, V. S. Nucleic Acid Research 2009, 37, D436–D442.

(4) Roos, W. H.; Bruinsma, R.; Wuite, G. J. L. Nature Physics 2010, 6, 733–743.

(5) Roos, W. H.; Wuite, G. J. L. Adv. Mater 2009, 21, 1187–1192.

(6) Ayton, G.; Voth, G. A. Biophys. J. 2010, 99, 2757–2765.

(7) Miao, Y.; Johnson, J. E.; Ortoleva, P. J. J. Phys. Chem. B 2010, 114, 11181–11195.

(8) Ode, H.; Nakashima, M.; Kitamura, S.; Sugiura, W.; Sato, H. Front. Microbiol. 2012, 3, 1–9.

(9) Zhao, G.; Perilla, J. R.; Yufenyuy, E. L.; Meng, X.; Chen, B.; Ning, J.; Ahn, J.; Gronenborn, A. M.; Schulten, K.; Aiken, C.; Zhang, P. Nature 2013, 497, 643–646.

(10) Bancroft, J. B.; Hiebert, E. Virology 1967, 32, 354–356.

(11) Arkhipov, A. Biophys. J. 2009, 97, 2061–2069.

(12) Globisch, C.; Krishnamani, V.; Deserno, M.; Peter, C. PLoS ONE 2013, 8, e60582.

(13) Kononova, O.; Snijder, J.; Brasch, M.; Cornelissen, J.; Dima, R. I.; Marx, K. A.; Wuite, G. J. L.; Roos, W. H.; Barsegov, V. Biophys. J. 2013, 105, 1893–1903.

(14) Sanner, M. F.; Olson, A. J. Biopolymers 1996, 38, 305–320.

(15) Chen, M.; Lu, B. J. Chem. Theory Comput. 2011, 7, 203–212.

(16) Krone, M.; Stone, J.; Ertl, T.; Schulten, K. Fast Visualization of Gaussian Density Surfaces for Molecular Dynamics and Particle System Trajectories. EuroVis - Short Papers. 2012.

(17) Si, H. ACM Trans. Math. Softw. 2015, 41, 1–36.

(18) Aggarwal, A.; Chen, J.-S.; Klug, W. S. Comput. Model. Eng. Sci. 2014, 98, 69–99.

(19) Carrillo-Tripp, M.; Brooks, C. L.; Reddy, V. S. PROTEINS: Structure, Function, and Bioinf. 2008, 73, 644–655.

(20) Berman, H. M.; Westbrook, J.; Feng, Z.; Gilliland, G.; Bhat, T. N.; Weissig, H.; Shindyalov, I. N.; Bourne, P. E. Nucleic Acids Research 2000, 28, 235.

(21) Housecroft, C.; Sharpe, A. G. Inorganic Chemistry; Pearson (4th ed.), 2012; pp 1013–1014.

(22) Zienkiewicz, O. C.; Morgan, K. Finite elements and approximation.

(23) Gibbons, M. M.; Klug, W. S. Phys. Rev. E 2007, 75, 031901.

(24) Gibbons, M. M.; Klug, W. S. Biophys. J. 2008, 95, 3640–3649.

(25) Roos, W.; Gibbons, M.; Arkhipov, A.; Uetrecht, C.; Watts, N.; Wingfield, P.; Steven, A.; Heck, A.; Schulten, K.; Klug, W.; Wuite, G. Biophys. J. 2010, 99, 1175–1181.

(26) Michel, J. P.; Ivanovska, I. L.; Gibbons, M. M.; Klug, W. S.; Knobler, C. M.; Wuite, G. J. L.; Schmidt, C. F. PNAS 2006, 103, 6184–6189.

(27) Uetrecht, C.; Versluis, C.; Watts, N. R.; Roos, W. H.; Wuite, G. J. L.; Wingfield, P. T.; Steven, A. C.; Heck, A. J. R. PNAS 2008, 105, 9216–9220.

(28) Hernando-Perez, M.; Pascual, E.; Aznar, M.; Ionel, A.; Caston, J. R.; Luque, A.; Carrascosa, J. L.; Reguera, D.; de Pablo, P. J. Nanoscale 2014, 6, 2702–2709.

(29) Roos, W. H.; Gertsman, I.; May, E. R.; III, C. L. B.; Johnson, J. E.; Wuite, G. J. L. PNAS 2012, 109, 2342–2347.

(30) Vargas-Felix, J. M.; Botello-Rionda, S. Acta Universitaria 2012, 22, 14–24.

(31) Aggarwal, A.; May, E. R.; Brooks, C. L.; Klug, W. S. Phys. Rev. E 2016, 93, 012417.

